# Indoleamine 2, 3-dioxygenase 1 mediated alterations in the functionality of immune cells, decipher the pregnancy outcomes in crossbred dairy cows

**DOI:** 10.1101/2022.08.01.502424

**Authors:** Sunil Kumar Mohapatra, Dheeraj Chaudhary, Bibhudatta S. K. Panda, Yallappa M. Somagond, Aarti Kamboj, Rajeev Kapila, Ajay Kumar Dang

**Affiliations:** Department of Animal Biochemistry, ICAR-National Dairy Research Institute, Karnal, Haryana 132001, India; Lactation and Immuno-Physiology Laboratory, ICAR-National Dairy Research Institute, Karnal, Haryana 132001, India

**Keywords:** IDO1, Neutrophil functionality, PBMCs functionality, pregnant cows

## Abstract

Pregnancy establishment in bovines requires maternal immune cell modulation. Present study investigated possible role of immunosuppressive indolamine-2, 3-dioxygenase 1 (IDO1) enzyme in the alteration of neutrophil (NEUT) and peripheral blood mononuclear cells (PBMCs) functionality of crossbred cows. Blood was collected from non-pregnant (NP) and pregnant (P) cows, followed by isolation of NEUT and PBMCs. Plasma pro-inflammatory (IFNγ and TNFα) and anti-inflammatory cytokines (IL-4 and IL-10) were estimated by ELISA and analysis of IDO1 gene in NEUT and PBMCs by RT-qPCR. Neutrophil functionality was assessed by chemotaxis, measuring activity of myeloperoxidase and β-D glucuronidase enzyme and evaluating nitric oxide production. Changes in PBMCs functionality was determined by transcriptional expression of pro-inflammatory (IFNγ, TNFα) and anti-inflammatory cytokine (IL-4, IL-10, TGFβ1) genes. Significantly elevated (P < 0.05) anti-inflammatory cytokines, increased IDO1 expression, reduced NEUT velocity, MPO activity and NO production observed only in P cows. Significantly higher (P < 0.05) expression of anti-inflammatory cytokines and TNFα genes were observed in PBMCs. Study highlights possible role of IDO1 in modulating the immune cell and cytokine activity during early pregnancy and may be targeted as early pregnancy biomarkers.

## 1. Introduction

The period of pregnancy exhibits a unique and idiosyncratic way of immune modulation for the dam and is an orchestrated event that requires the balance between immune activation and suppression. Mechanisms such as IFNτ mediated maternal recognition of pregnancy (MRP) (Raheem, 2017), modulation of immune cells function via different hormones such as progesterone and chorionic gonadotropin, decreased expression of major histocompatibility proteins by the trophoblast (Oliveira and Hansen, 2008), conceptus mediated modulation of immune-related genes, a critical balance between pro-inflammatory i.e., T-helper 1 cytokines (Th1 cytokines) and anti-inflammatory i.e., T-helper 2 cytokines (Th2 cytokines) pathway (Clark et al., 2005; Morelli *et al*., 2015), macrophages recruitment and plasticity, immune-modulation via secreted fetal DNA into maternal circulation (Oliveira et al., 2012) and many other mechanisms which have been studied in detail to unravel the mechanisms behind maternal immune-modulation during the early phase of bovine pregnancy. All such mechanisms of maternal immune-modulation can be understood by two different perspectives; one may be due to the direct effect of the major immune cells such as neutrophils (first line of defense), lymphocytes and macrophages during pregnancy establishment and second is the role of various factors that indirectly affects the function of different immune cells.

Effect of immune cells on maternal immune modulation broadly includes pregnancy-associated uterine remodeling specifically by the uterine natural killer cells (uNKs), macrophages, and T cells, vascular remodeling by the uterine mast cells (uMCs), shifting of cytokines towards anti-inflammatory pathway, changes in the placental morphology, expression of immune-related genes by the endometrial cells, etc., (Zenclussen and Hämmerling, 2015; Meyer and Zenclussen, 2020) whereas, the factors affecting the function of the immune cells during the early phase of bovine pregnancy involve various hormones, cytokines, enzymes, fetal DNA, etc. (Bonney, 2016). Both the process goes hand-in-hand to prevent rejection of the semi-allogeneic fetus so as to finally achieve the pregnancy success. However, the exact underlying mechanism behind the precise and controlled immune-activation during early pregnancy establishment in bovines still remains elusive. Circulating immune cells migrate from peripheral circulation towards the feto-maternal interface and positively contribute towards maternal tissue remodeling and embryo-maternal cross-talk around the implantation period which is regulated under the influence of endocrine system (Fujiwara, 2009). Prevention of semi-allogeneic fetus from undergoing rejection is taken care of by the modulation of various maternal immune cells which drive the cytokine balance towards the anti-inflammatory Th2 pathway (Morelli et al., 2015). This immune tolerance character exhibited by the immune cells might be triggered via several molecules like hormones, cytokines, enzymes etc., (Bonney, 2016). Out of several factors modulating the immune cells, one important molecule is indoleamine 2, 3-dioxygenase (IDO) which is expressed in the immune cells (neutrophils and lymphocytes) present at both the feto-maternal interface as well as at the systemic level under the influence of IFNτ and/or IFNγ (Mohapatra et al., 2020). Groebner et al. (2011) have also highlighted the importance of increased IDO expression and activity in bovine pregnancy depicting its pivotal role in protecting the semi-allogeneic conceptus from maternal rejection. On the other hand, some reports also suggest that a dysfunctional and abnormal IDO expression is associated with several pathological pregnancies like pre-term labor, preeclampsia, recurrent spontaneous abortion and fetal growth restriction in human (Chang et al., 2018). IDO is an immunosuppressive enzyme that catabolizes the first and rate-limiting step of tryptophan catabolism. It depletes the available tryptophan present in the microenvironment and inhibits the growth and differentiation of alloreactive T-cells thus, skewing the differentiation of T-cells into Th2 cells (Munn et al., 1998). However, the significance of IDO expression in altering the biochemical functionality of immune cells has not been elucidated during the establishment of bovine pregnancy.

Despite thorough knowledge regarding the various mechanism of immune-modulation during early bovine pregnancy, the fundamental changes occurring in terms of the functionality of immune cells have not been addressed in bovines. Some studies have focused on the changes in number and types of immune cells (Oliveira and Hansen, 2008), whereas, few other studies have highlighted the changes at the transcriptomics and proteomics level in the immune cells during the early phase of pregnancy (Panda et al., 2020). However, the cellular, molecular and biochemical changes occurring in the immune cells (neutrophils and lymphocytes) and the mechanism behind such changes during the early phase of pregnancy in bovines still remains elusive. Therefore, the present novel study was undertaken to explore the major cellular, molecular and biochemical changes occurring in neutrophils and lymphocytes during the early phase of pregnancy vis-a-vis the significance of IDO expression behind such functionality changes of the immune cells. This study may be useful for establishing new ways of medical intervention to deal with the losses during pregnancy failure.

## 2. Results

### 2.1. Plasma levels of anti-inflammatory and pro-inflammatory cytokines in pregnant and non-pregnant cows

The plasma levels of anti-inflammatory cytokines (IL-4 and IL-10) and pro-inflammatory cytokines (IFNγ and TNFα) in non-pregnant and pregnant cows have been shown in Fig. 1A,B. The plasma concentration of IL-4 was observed to be remarkably high (P ≤ 0.001) in pregnant cows in contrast to the non-pregnant cows. A similar trend was also observed for IL-10 cytokine which increased tremendously (P ≤ 0.001) in pregnant cows as compared to the non-pregnant cows. However, the levels of both pro-inflammatory cytokines IFNγ and TNFα did not differ significantly between non-pregnant and pregnant cows.

**Fig. 1.**
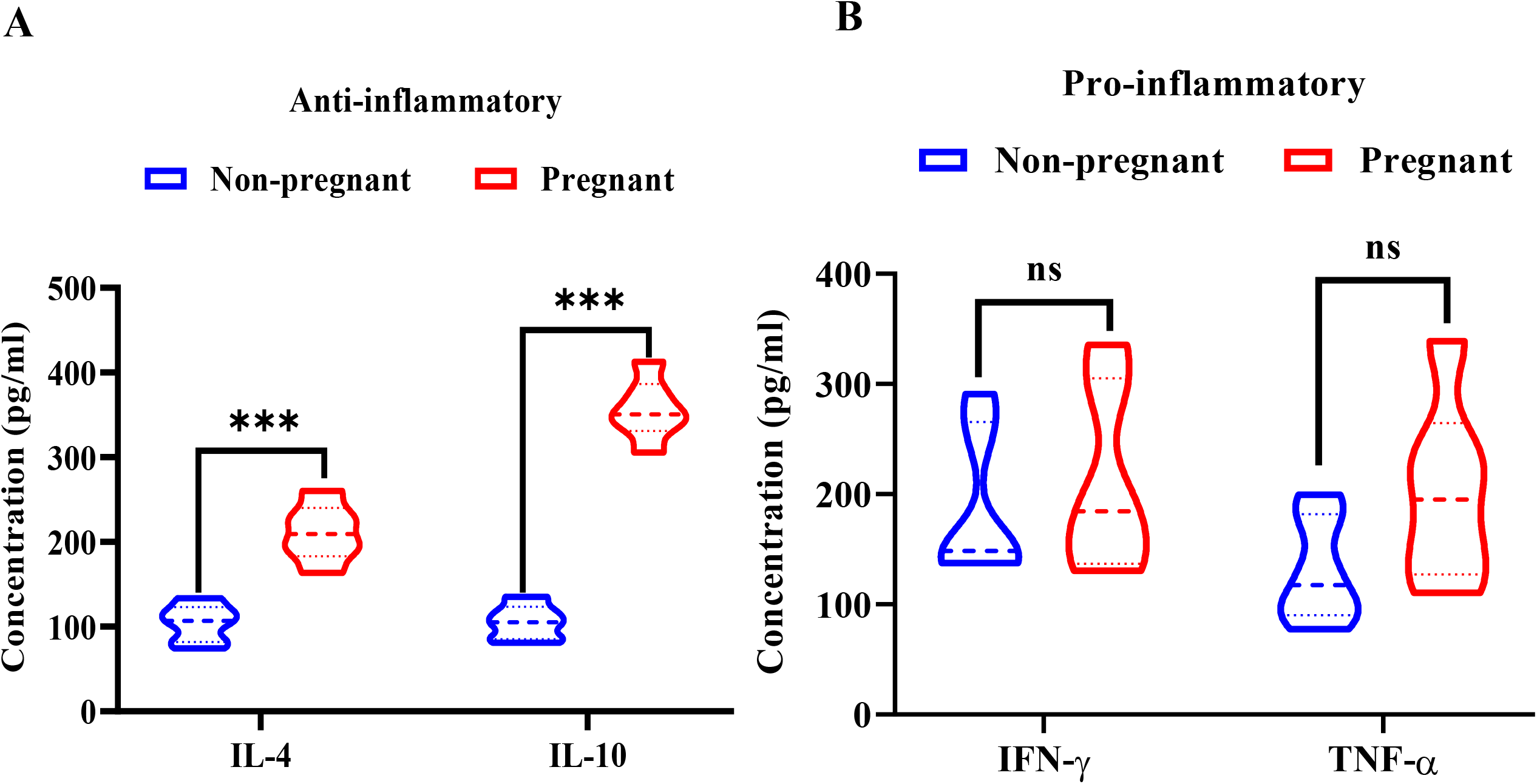
Concentration (pg/ml) of anti-inflammatory (IL-4 and IL-10) and pro-inflammatory (IFNγ and TNFα) cytokines in blood plasma of non-pregnant and pregnant cows. (A) Anti-inflammatory IL-4 and IL-10 cytokines in non-pregnant (blue, n=6) and pregnant (red, n=6) cows. (B) Pro-inflammatory IFNγ and TNFα cytokines in non-pregnant (blue, n=6) and pregnant (red, n=6) cows. All graphs are represented as mean and standard error of the mean (s.e.m.). Triple asterisks (***) represents a significant difference of P < 0.001 between the groups and ns represents a non-significant difference between the groups. Images are representative of three independent experiments.

### 2.2. Relative mRNA expression of indoleamine-2,3-dioxygenase1 (IDO1) in neutrophils and PBMCs of non-pregnant and pregnant cows

The mRNA expression of IDO1 gene was found to be significantly elevated in both neutrophils and PBMCs of pregnant cows as compared to the non-pregnant cows (Fig. 2). However, the fold change of IDO was higher in the case of PBMCs than in neutrophils. The fold change difference of IDO1 expression was observed to be 12 folds in PBMCs and 4 folds in neutrophils of the pregnant cows as compared to their respective non-pregnant counterparts.

**Fig. 2.**
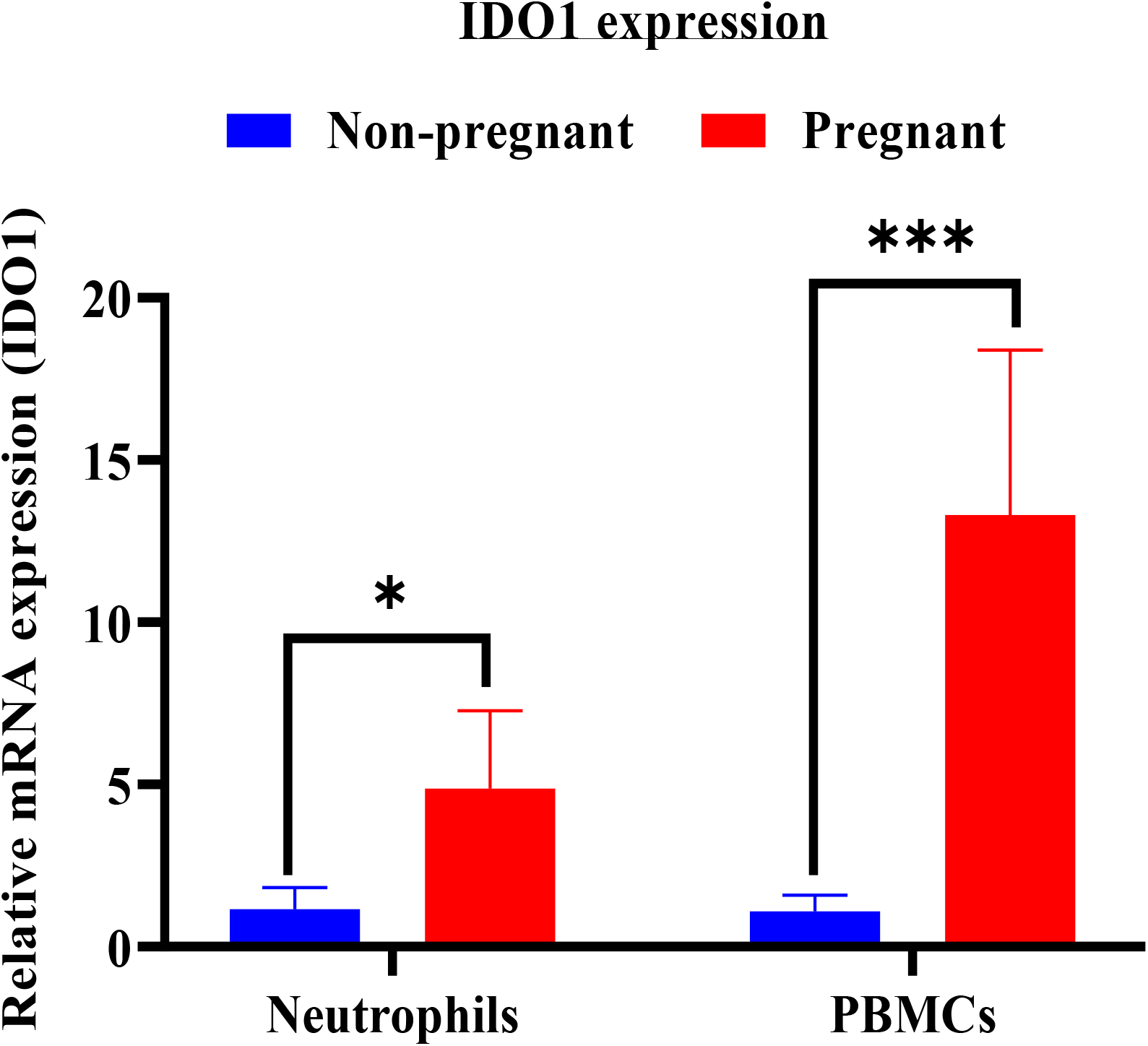
Relative mRNA expression of *IDO1* gene in neutrophils and PBMCs of non-pregnant and pregnant cows. Non-pregnant cows represented in blue (n=6) and pregnant cows represented in red (n=6). Graph is represented as mean and standard error of the mean (s.e.m.). Single asterisk (*) represents a significant difference of P < 0.05 between neutrophils of pregnant and non-pregnant groups. Triple asterisks (***) represents a significant difference of P < 0.001 between PBMCs of pregnant and non-pregnant groups. Images are representative of three independent experiments.

### 2.3. Evaluation of key neutrophil functions

The functionality of neutrophils in the test animals was assessed by chemotaxis (velocity, distance, displacement and directionality), measuring activity of marker enzymes (myeloperoxidase and β-D glucuronidase) and nitric oxide production.

The neutrophils from pregnant cows were observed to be less active as compared to the neutrophils of the non-pregnant cows. Less number of neutrophils from pregnant cows was spotted to be migrated towards the chemoattractant in contrast to the neutrophils from the non-pregnant cows (Fig. 3A,B).

**Fig. 3.**
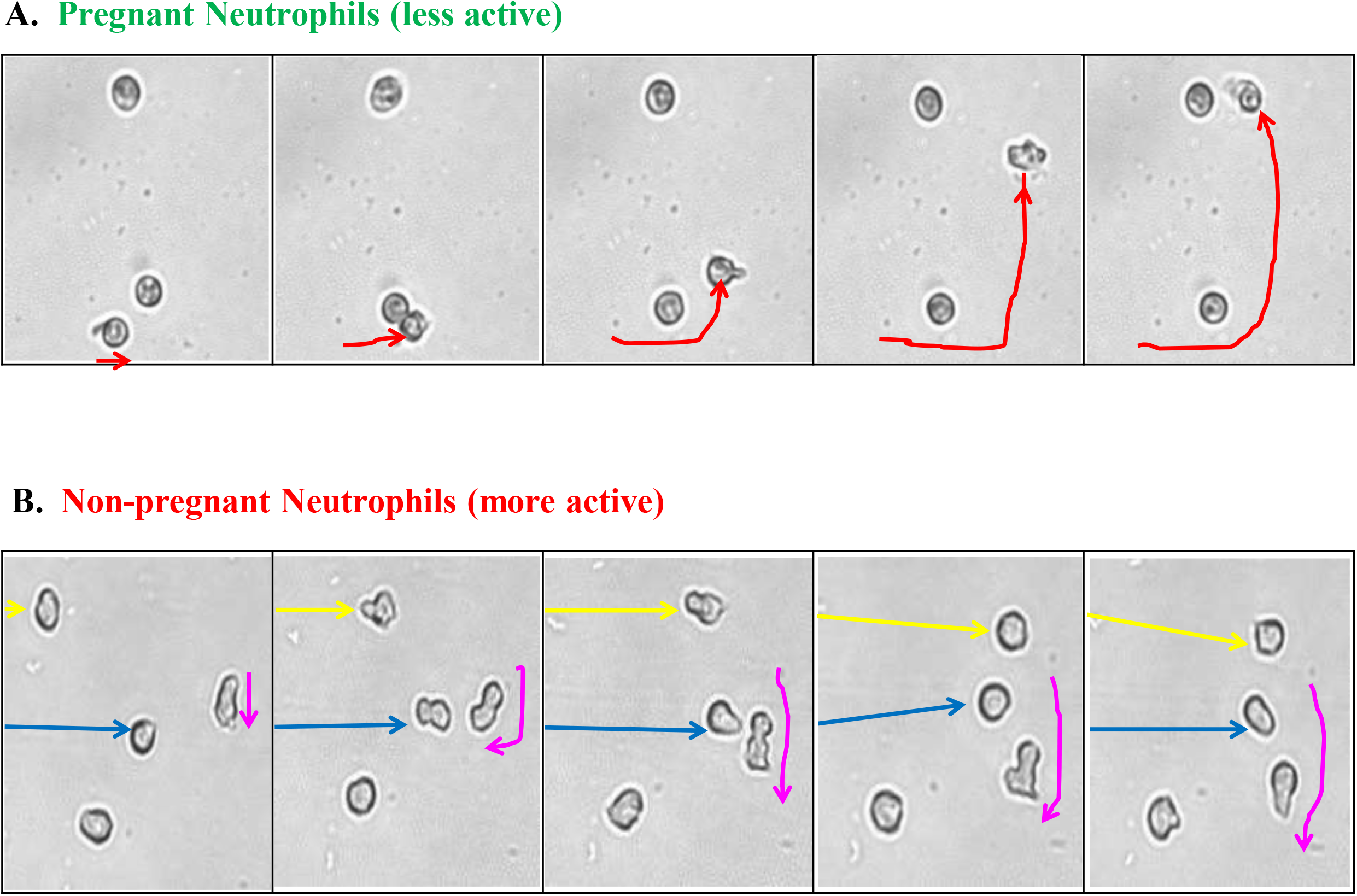
Functionality of neutrophils assessed by chemotaxis. (A) Representative migration image of pregnant neutrophils assessed by time-lapse microscopy. Red arrow represents the migration of pregnant neutrophil. Length of arrow indicates the distance covered by the neutrophils. (B) Representative migration image of non-pregnant neutrophils assessed by time-lapse microscopy. Yellow, pink and blue arrow represents the migration of non-pregnant neutrophils. Each coloured arrow represents individual neutrophil migration. Length of arrow indicates the distance covered by the neutrophils. Images are representative of three independent experiments.

The velocity, total distance travelled, displacement and directionality of the neutrophils from non-pregnant and pregnant cows have been illustrated in Fig. 4A,B,C,D respectively. The velocity of neutrophils from pregnant animals was observed to be significantly lower (P ≤ 0.05) than the neutrophils of non-pregnant ones. However, the other factors such as distance, displacement and directionality of neutrophils did not show any significant difference statistically and remained similar in both non-pregnant and pregnant cows.

**Fig. 4.**
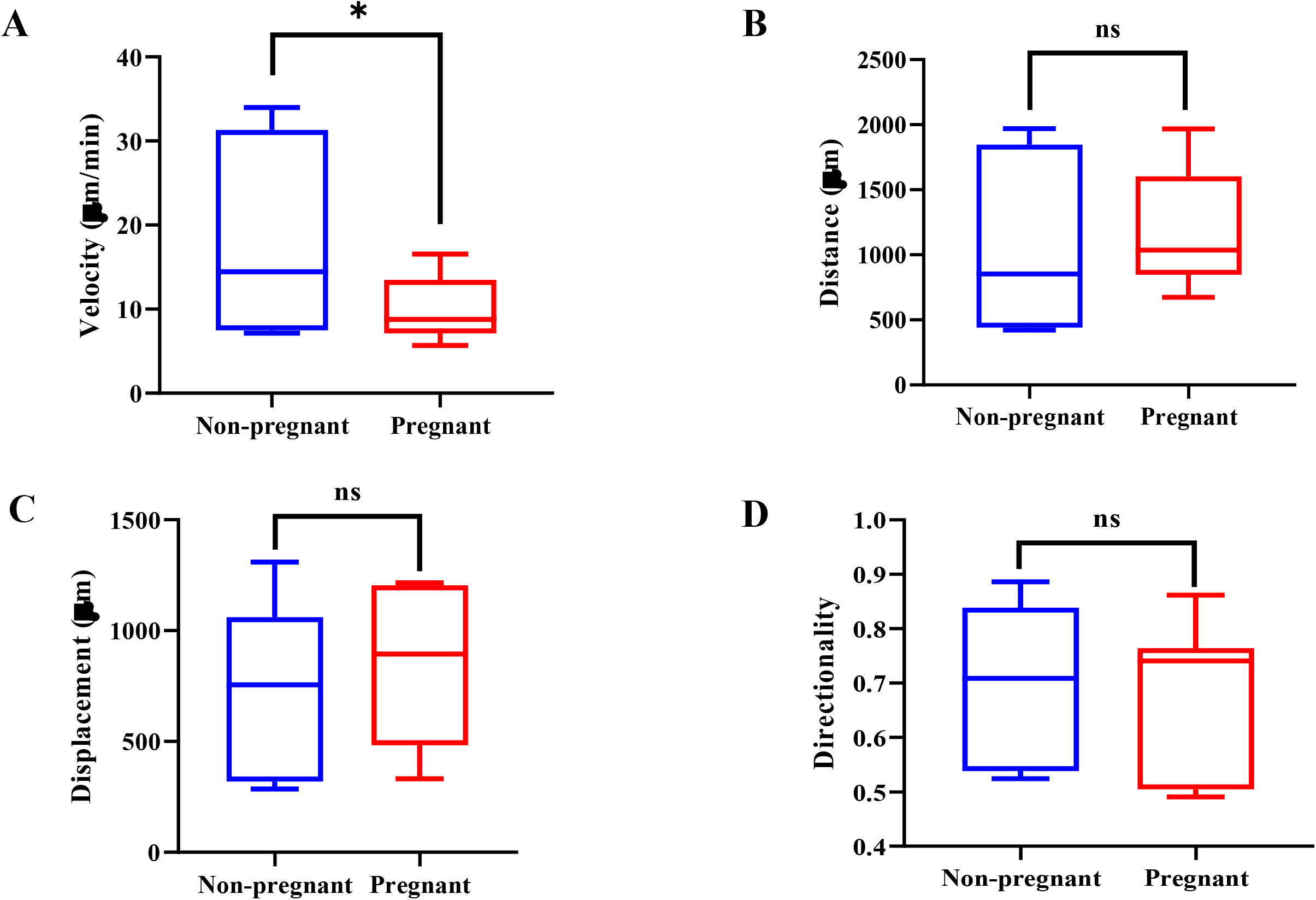
Neutrophil functionality in terms of velocity, total distance travelled, displacement and directionality of the neutrophils from non-pregnant and pregnant cows. (A) Velocity of neutrophils obtained from non-pregnant (blue, n=6) and pregnant (red, n=6) cows. (B) Total distance travelled by the neutrophils obtained from non-pregnant (blue, n=6) and pregnant (red, n=6) cows. (C) Displacement of neutrophils obtained from non-pregnant (blue, n=6) and pregnant (red, n=6) cows. (D) Directionality of neutrophils obtained from non-pregnant (blue, n=6) and pregnant (red, n=6) cows. All graphs are represented as mean and standard error of the mean (s.e.m.). Single asterisk (*) represents a significant difference of P < 0.05 between neutrophils of pregnant and non-pregnant groups. ns represents a non-significant difference between the groups. Images are representative of three independent experiments. All analysis was performed using Image-J (Chemotaxis tool) software.

The myeloperoxidase (MPO) activity of neutrophils from non-pregnant and pregnant cows has been portrayed in Fig. 5A. In pregnant cows, the MPO activity of neutrophils was found to be significantly reduced (P ≤ 0.05) as compared to the MPO activity of neutrophils in non-pregnant cows.

**Fig. 5.**
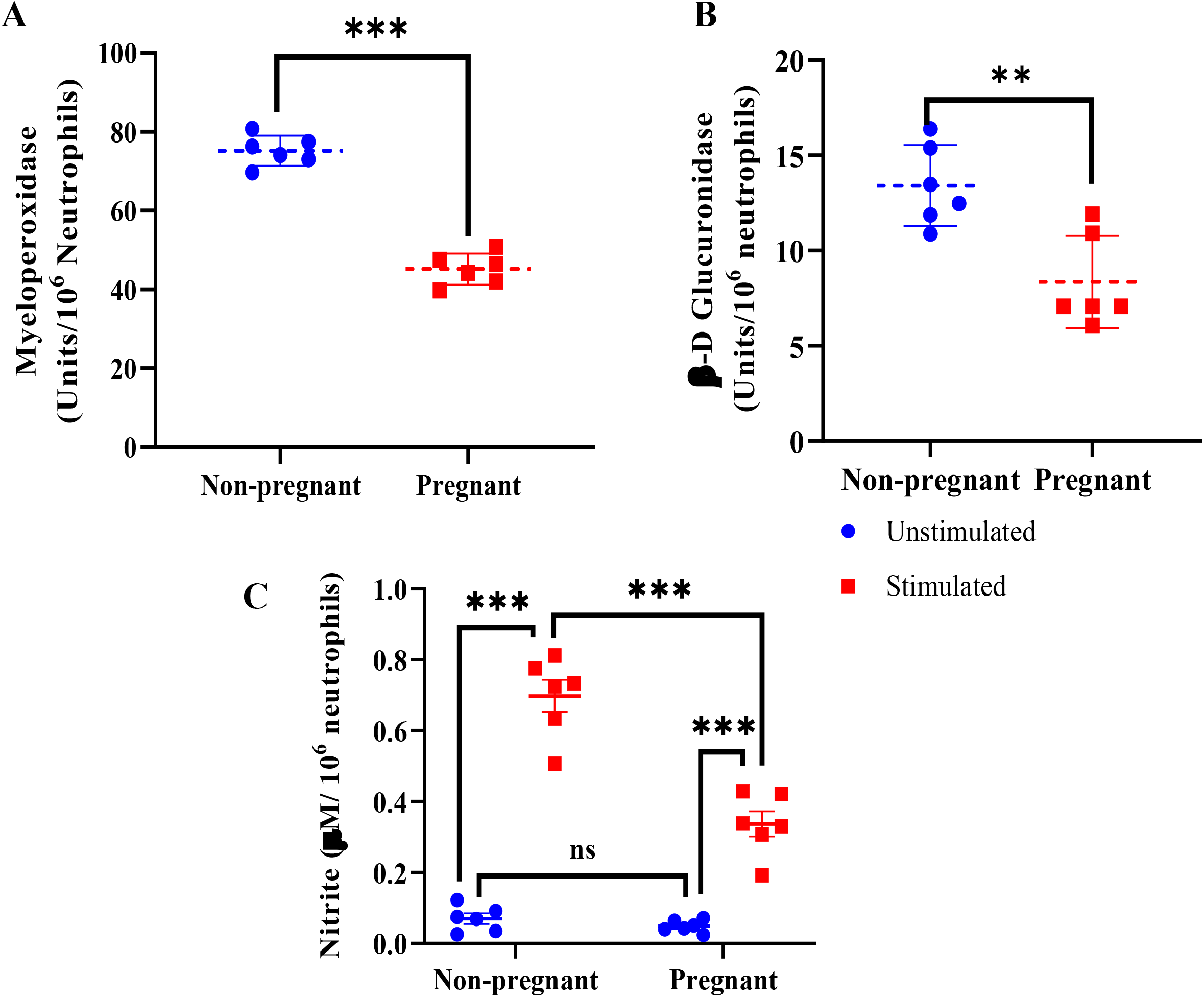
Neutrophil functionality in terms of neutrophil degranulation. (A) Myeloperoxidase activity in neutrophils of non-pregnant (blue, n=6) and pregnant (red, n=6) cows. (B) β-glucuronidase activity in neutrophils of non-pregnant (blue, n=6) and pregnant (red, n=6) cows. (C) Nitric oxide assay in neutrophils of non-pregnant (blue, n=6) and pregnant (red, n=6) cows. All graphs are represented as mean and standard error of the mean (s.e.m.). Double asterisks (**) represents a significant difference of P < 0.01 between neutrophils of pregnant and non-pregnant groups. Triple asterisks (***) represents a significant difference of P < 0.001 between the groups. ns represents a non-significant difference between the groups. Images are representative of three independent experiments.

The β-glucuronidase activity of neutrophils from non-pregnant and pregnant cows has been presented in Fig. 5B. The activity of β-glucuronidase in the neutrophils of pregnant cows significantly lowered (P ≤ 0.05) as compared to the β-glucuronidase activity in neutrophils of non-pregnant cows.

The nitrite oxide level of neutrophils from non-pregnant and pregnant cows has been indicated in Fig. 5C. The analysis of nitrite oxide levels revealed a significant reduction (P ≤ 0.05) in the LPS stimulated neutrophils of pregnant cows as compared to the LPS stimulated neutrophils of non-pregnant cows. However, no significant difference in the nitrite levels was observed between the unstimulated neutrophils of pregnant and non-pregnant cows.

### 2.4. Evaluation of PBMCs functionality

The release of cytokines by the PBMCs is regarded among their key functional parameters which has been determined through relative mRNA expression of canonical pro-inflammatory (IFNγ and TNFα) and anti-inflammatory (IL-4, IL-10 and TGFβ) genes.

The relative mRNA expressions of anti-inflammatory cytokines (IL-4, IL-10 and TGFβ1) genes in PBMCs of non-pregnant and pregnant cows have been depicted in Fig. 6A. In case of pregnant cows, the relative mRNA expression of IL-4, IL-10 and TGFβ1 genes were significantly (P ≤ 0.05) up regulated as compared to their non-pregnant counterparts. The expression levels of IL-4 and IL-10 genes were observed to be around 20 folds higher, whereas that of TGFβ1 gene was around 4 fold higher in case of pregnant cows.

**Fig. 6.**
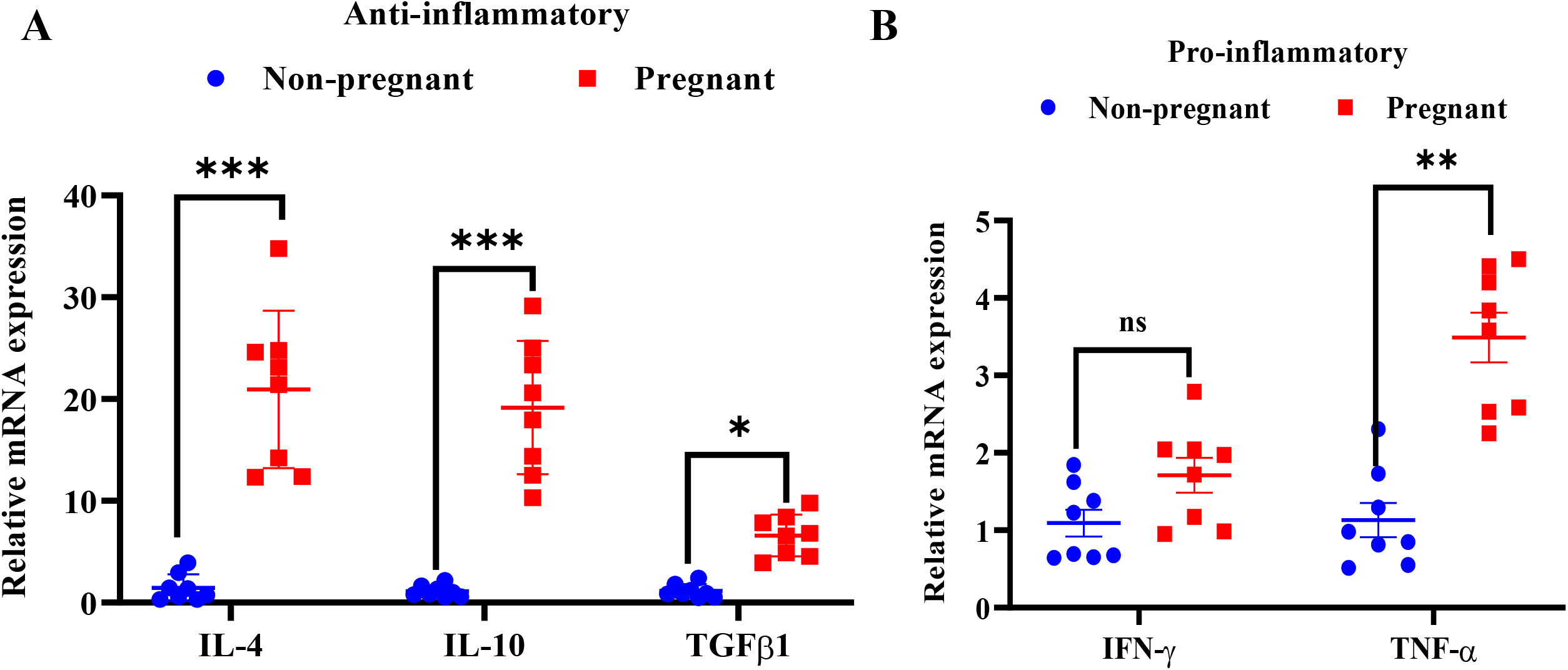
PBMCs functionality in terms of cytokine expression. (A) Relative mRNA expression of anti-inflammatory cytokines (IL-4, IL-10 and TGFβ1) in PBMCs of non-pregnant (blue, n=6) and pregnant (red, n=6) cows. (B) Relative mRNA expression of pro-inflammatory cytokines (IFNγ and TNFα) in PBMCs of non-pregnant (blue, n=6) and pregnant (red, n=6) cows. Single asterisk (*) represents a significant difference of P < 0.05 between neutrophils of pregnant and non-pregnant groups. Double asterisks (*) represents a significant difference of P < 0.01 between neutrophils of pregnant and non-pregnant groups. Triple asterisks (***) represents a significant difference of P < 0.001 between the groups. Images are representative of three independent experiments.

The relative mRNA expressions of pro-inflammatory (IFNγ and TNFα) cytokine genes in PBMCs of non-pregnant and pregnant cows have been shown in Fig. 6B. The relative abundance of TNFα mRNA was observed to be significantly more (P ≤ 0.05) in pregnant cows as compared to the non-pregnant cows. However, the expression level of the IFNγ gene did not change much statistically between pregnant and non-pregnant cows.

## 3. Discussion

The exact mechanism behind maternal immune-modulation during early pregnancy establishment in cattle is still unexplored. A plethora of literatures emphasize the regulation and importance of immune cells at the juncture of maternal-fetal interface for pregnancy establishment and any imbalance in the harmony causes adverse pregnancy outcomes (Yang et al., 2019; Than et al., 2019). Moreover, several tolerogenic factors are known to modulate the immune cells towards a secure pregnancy. One of such factor is indoleamine-2, 3-dioxygenase (IDO) which needs to be critically explored in the context of bovine pregnancy. Therefore, in the present study, we investigated and compared the differential functionality of immune cells (particularly neutrophils and PBMCs) in pregnant and non-pregnant cows in relation to IDO1 expression.

IDO1 is the first and rate-limiting enzyme in tryptophan catabolism and acts via the kynurenine pathway to convert tryptophan into N-formyl-kynurenine and other biologically active compounds (Munn et al., 1998). IDO1 causes immunosuppression through utilization of the available tryptophan present in the microenvironment and thereby causes starvation of the neighboring maternal T-cell for this essential amino acid. Subsequently, the deficiency of tryptophan blocks the progression of cell cycle in activated alloreactive T-cells which leads to T-cell arrest at mid-G1 phase of the cell cycle and the cells undergo apoptosis (Lee et al., 2002). This process leads to immunosuppression due to the inhibition of locally activated T-cell proliferation. Few studies in the context of human pregnancy has revealed that higher IDO1 expression in trophoblast cells, chorionic vascular endothelial cells, stromal cells and immune cells of deciduata, peripheral blood immune cells is the hallmark of successful pregnancy whereas, dysregulated IDO1 expression leads to pregnancy losses (Zong et al., 2016; Chang et al., 2018). However, in the context of bovine pregnancy, the role of IDO1 is still largely unknown. Few researchers have highlighted that in bovines, the resident immune and non-immune cells of the endometrium display greater IDO1 expression during pregnancy (Kamat et al., 2016). In our current study, more IDO1 expression in neutrophils and PBMCs were associated with successful pregnancy and it is in accordance with our earlier study where we recorded identical findings in pregnant cows (Mohapatra et al., 2020). The probable reason may be because IDO1 metabolizes L-tryptophan present in the micro environment and curbs the proliferation of alloreactive T cells which might be detrimental for the semi-allogenic embryo (Munn et al., 1998). Moreover, shreds of evidence support that tolerogenicity can also be achieved indirectly due to the IDO1 mediated Th1-Th2 paradigm shift in different immune cells (Xu et al., 2008; Krupa and Kowalska, 2021).

A variety of immune cells like natural killer cells, macrophages, T cells, mast cells, etc., takes part in the modulation of maternal immune system during pregnancy. One such important immune cell is neutrophils which are short-lived granulocytic leucocytes having the characteristic multi-lobed nucleus and different cytoplasmic granules (both acidic and basic granules). Traditionally, they are regarded as the first line of cellular defense against infection. However, in the context of pregnancy their contribution towards successful conception, implantation, pregnancy maintenance and parturition is paramount whereas, impaired neutrophil functionality is catastrophic for it (Hahn et al., 2012; Alhussein and Dang, 2019). Thus, in our study, we framed to decipher the functionality changes that occur in the bovine neutrophils during pregnant and non-pregnant conditions. Owing to an alien intrusion, neutrophils rapidly migrate towards the site of inflammation. It is mediated by different chemo-attractants which enable the directional migration of neutrophils along the concentration gradient through the process of chemotaxis (Metzemaekers et al., 2020). Chemotaxis is central to combat against the foreign antigen for which speed and directionality are the measures for it (Hoang et al., 2013). Krause et al. (1987) observed a depressed chemotaxis in the neutrophils obtained from the pregnant females. Similarly, in our study, we also observed a significant reduction in the velocity of pregnant bovine neutrophils. The probable reason for the decreased responsiveness of pregnant neutrophils might be due to the increased kynurenine concentration in the plasma because of up-regulated IDO1 activity and it is well known that kynurenine is a potent inhibitor of neutrophil chemotaxis (Loughman et al., 2016).

In order to eliminate intruders, the activated neutrophils release myeloperoxidase (MPO) i.e. a pro-inflammatory enzyme stored in the azurophilic granules, into both the phagolysosomal compartment as well as to the extracellular environment. Neutrophils with the help of MPO exhibit its microbicidal activity through the generation of reactive oxygen species (ROS). MPO catalyzes the hypochlorous acid formation from the H_2_O_2,_ which is responsible for the microbicidal activity of neutrophils during the active inflammatory state. Therefore, assessing the MPO activity in neutrophils will indicate its inflammatory status (Pulli et al., 2013). In our study, we noticed lower MPO activity in the pregnant neutrophils which might be associated with low inflammatory condition and thus maintenance of the pregnancy. Elevated MPO activity has been observed as detrimental for successful pregnancy outcome (Rocha-Penha et al., 2017). It has also been noted that higher MPO concentration in neutrophils is correlated with the embryonic mortality in bovines (Panda et al., 2020).

Apart from ROS production, neutrophils also produce nitric oxide during inflammatory conditions which can be assessed indirectly by measuring the accumulation of the stable end product nitrite using a modified Griess reaction method Kasai et al., (1995). There are very limited literatures available in the public domain depicting the importance of nitric oxide production during pregnancy. However, some studies in humans revealed that, the nitric oxide synthesis in neutrophils was more in pregnant women as compared to the non-pregnant women (Tsukimori et al., 2006). In stark contrast to this, we recorded a low amount of nitric oxide in neutrophils obtained from the pregnant cows. Present works of literature available have a huge gap in elucidating the significance of nitric oxide in the context of pregnancy and further research is required to explore this untapped area.

Another important molecule stored in the azurophilic granules of the neutrophils is the β-glucuronidase enzyme. It is mainly a lysosomal hydrolase enzyme which is widely distributed in the mammalian tissue. β-glucuronidase enzyme is a potent inflammatory agent involved during various inflammatory conditions. This enzyme has been widely targeted as the agent for anticancer chemotherapy (Boyland et al., 1957), treatment of neonatal jaundice (Gourley, 2002), etc. However, the role of β-glucuronidase in the context of bovine pregnancy has been poorly elucidated and very limited literatures are available depicting its significant role during pregnancy. Amagase et al. (1997) reported no significant difference in the β-glucuronidase activity in the neutrophils of the non-pregnant and the normal pregnant women. However, in contrast to this, we noted significantly reduced level of β-glucuronidase activity in the neutrophils obtained from the pregnant cows. As β-glucuronidase is mainly involved to provoke the inflammatory conditions, therefore, its increased activity during pregnancy might be detrimental for the semi-allogeneic fetus and this might be the plausible cause for the reduction of its activity during bovine pregnancy. Thus, in order to elucidate the actual depth of the involvement of β-glucuronidase enzyme in the context of pregnancy, extensive and rigorous research needs to be conducted.

Apart from the involvement of neutrophils and its enzymes in pregnancy, another crucial aspect which needs to be balanced during early pregnancy establishment is the delicate harmony between pro-inflammatory (Th1) and anti-inflammatory (Th2) cytokines which is needed to increase the maternal tolerance towards the semi-allogeneic embryo (Wegmann et al., 1993). Any disturbance in the Th1:Th2 cytokine balance will lead to fetal death and failure in pregnancy (Sykes et al., 2012). During the time of implantation the immunosuppressive action of IDO1 abrogates the production of pro-inflammatory cytokines by suppressing the Th1 cell population and promotes Th2 cells like T-reg cells to produce more anti-inflammatory cytokines for the embryo survival (Xu et al., 2008). The production of these cytokines by Th1 and Th2 cells are regarded as the key functional parameters of the lymphocytes. Therefore, in order to determine the change in the functionality of PBMCs during early pregnancy, their ability of cytokine synthesis and secretion needs to be assessed (Navas et al., 2019; Chen et al., 2020). For this, we studied the mRNA abundances of different Th2 cytokine genes (IL-4, IL-10 and TGF-β1) and Th1 cytokine genes (IFN*-*γ and TNFα) in PBMCs of pregnant and non-pregnant cows in correlation with the IDO1 expression. Further, we also validated it by measuring their respective plasma levels through ELISA. Our observations delineated significantly higher (P < 0.05) relative mRNA expression levels of various anti-inflammatory cytokines (IL-4, IL-10 and TGF-β1) genes in pregnant cows compared to the non-pregnant cows.

Interleukin-4 (IL-4), an anti-inflammatory cytokine is known to promote the differentiation of naive helper T cells (Th0 cell) towards Th2 cell and subsequently these Th2 cells produce additional IL-4 in a positive feedback loop (Gool, 1999). IL-4 decreases the differentiation of Th1 cells and is a key regulator of humoral and adaptive immunity (Sokol et al., 2008). In our study, we observed significantly increased (P < 0.05) real-time expressions of IL-4 gene as well as high blood plasma concentration of IL-4 in the pregnant cows. As IL-4 is responsible for inhibiting the function and development of Th1 cells, therefore, it might provide an anti-inflammatory milieu for the developing embryo and this could be the possible explanation of getting relatively higher mRNA abundance as well as high blood plasma concentration of IL-4 in pregnant cows in comparison to the non-pregnant cows. Our results are in line with Manjari et al., 2018 and Yang et al., 2018 in which they documented the elevated blood plasma level of IL-4 and higher mRNA as well as protein expression in PBMCs of pregnant cows around the peri-implantation period. Nonetheless, IL-4 is a negative regulator of IDO1 expression in human monocytes and this safeguards the animal from tryptophan metabolites having toxic potential and also from a state of excessive immune suppression (Musso et al., 1994); although this mechanism in the context of bovine pregnancy still needs to be explored.

Along with interleukin-4, interleukin-10 (IL-10) cytokine is also necessary for pregnancy (Chatterjee et al., 2014). IL-10 cytokine is a potent anti-inflammatory agent primarily produced by lymphocytes, monocytes and Th2 cells under the influence of systemic IFNτ (Shirasuna et al., 2011). In our study, we observed a significant increase (P < 0.05) in the *IL-10* mRNA expression as well as high plasma concentration of IL-10 in pregnant cows compared to the non-pregnant ones. Being an anti-inflammatory cytokine, IL-10 is considered to have multifaceted roles during pregnancy. According to Cheng and Sharma (2015), IL-10 suppresses the fetal-specific T cell proliferation and facilitates differentiation of lymphocytes subsets like Treg cells which favours the immune tolerant environment for the conceptus. Moreover, IL-10 is one of the potent regulators of IDO expression (Munn et al., 2002) and these might be the plausible causes of noticing abundant mRNA expression and plasma concentration in pregnant animals.

Besides interleukin-10, transforming growth factor beta 1 (TGF-β1) is another multifunctional cytokine that exhibits potential immuno-regulatory and anti-inflammatory properties for the survival and growth of the allograft. It dampens the dam’s inflammatory response to allow fetal growth and development (Figueiredo and Schumacher, 2010). The level of TGF-β1 has been reported to be higher in pregnant women, which revealed its potential regulatory function in fetal allograft survival in successful pregnancies (Ayatollahi et al., 2007). Further, it also aids in angiogenesis, Treg production, balance between M1-M2 macrophage polarization and regulation of the NK cell function (Yang et al., 2021). In our study, we reported significantly increased (P < 0.05) mRNA expression of *TGF-β1* in PBMCs of the pregnant cows. Elucidating its significance in the context of bovine pregnancy and *IDO1* expression, various researchers suggest that TGF-β1 works synergistically with IDO1 in the conversion of naive T cell into Treg cell via increasing kynurenine formation (Gandhi et al., 2010). TGF-β1 also up-regulates the IDO1 expression in the placenta of human (Wang et al., 2020) and this might be a cause of long term IDO1 expression and the resulting dominant Th2 cytokine status during pregnancy. Nevertheless, the up-regulation in the expression of these anti-inflammatory cytokine genes i.e., *IL-4, IL-10* and *TGF-β1* in pregnant cows might be necessary for the prevention of semi-allogeneic fetal rejection and thereby successful pregnancy establishment in cattle.

Up-regulation in the pro-inflammatory cytokines (Th1 cytokines) like interferon gamma (IFNγ) and Tumor Necrosis Factor Alpha (TNFα) are believed to be deleterious for the fetal survival during pregnancy establishment (Robertson et al., 2018). On a positive note, IFN-γ is a potent stimulator of IDO1 expressions at the feto-maternal interface during the first trimester of human pregnancy (Kudo et al., 2004). While on a negative note, aberrant expression and concentration of IFN-γ might lead to pathological pregnancy (Yockey and Iwasaki, 2018). IFN-γ prevents the Th2 response and favours the Th1 immune response by mediating the naive T cell differentiation into Th1 cells (Nakagome et al., 2009). According to Yang et al. (2018), PBMCs isolated from non-pregnant cows showed significantly higher (P < 0.05) expression of IFN-γ as compared to pregnant cows. But our study revealed no significant difference in the relative mRNA expression of IFN-γ between pregnant and non-pregnant cows and we are in agreement with the report of Vasudevan et al. (2017) that irrespective of pregnancy status or days there was no change in IFN-γ mRNA abundance in the uterus of bovine. Nonetheless, during the peri-implantation period in bovine, IFNτ, the only known agent of MRP is instrumental for IDO1 expression.

On the other hand, TNFα is a pro-inflammatory cytokine having pleiotropic effects during pregnancy. Elevated TNFα expression in human PBMCs has been observed to be associated with different forms of pregnancy complications (Azizieh and Raghupathy, 2015).

On contrary, higher concentration of TNFα assists in the process of implantation, decidualization and placentation during pregnancy (Ozbilgin et al., 2015). In our study, we observed significantly elevated (P < 0.05) levels of TNFα expression in the PBMCs of pregnant cows. Our results are in accordance with Sakumoto et al. (2014) who reported higher levels of TNFα mRNA and protein in the pregnant cows compared to the cyclic cows. The PBMCs derived TNFα modulates PGE2 production (luteotrophic) and reduces PGF2α secretion (luteolytic) during early phase of pregnancy as reported by Szostek et al. (2014). It also drives the vascularization of CL and hence angiogenesis (Fair et al., 2015). This may be the credible cause for the elevated TNFα expression in the pregnant cows which might be essential for the uterine function and conceptus development during early phase of pregnancy.

## 4. Materials and methods

### 4.1. Animal ethics approval and consent to participate

Clearance of the present study was taken from the Institutional Animal Ethics Committee (Approval No. 41-IAEC-18-32) of Indian Council of Agricultural Research (ICAR)-National Dairy Research Institute (NDRI) constituted as per the article 13 of the Committee for the Purpose of Control and Supervision of Experiments on Animals (CPCSEA) rules, laid down by the Government of India (Reg. No. 1705/GO/Re/SL/13/CPCSEA dated. 3/7/2013). All the ethical guidelines were followed during the course of the experiment.

### 4.2. Study site and selection of experimental cows

The present research was conducted at the Livestock Research Centre (LRC) of ICAR-National Dairy Research Institute (NDRI), Karnal, Haryana, India. This institute is located at 250 meters above sea level in the Indo-Gangetic plains, at 29°43′′ north latitude and 77°2′′ east longitude.

To investigate the role of IDO in alteration of the functionality of neutrophils and peripheral blood mononuclear cells (PBMCs) during early pregnancy, a total of 12 healthy, multiparous, Karan Fries (KF) crossbred (Holstein Friesian x Tharparkar) cows of third to fifth parity, aged between 5–7 years with regular cyclicity and good health condition were selected and divided into two groups viz; (i) Pregnant (N=6) and (ii) Non-pregnant (N=6). Cows which did not return to heat with higher plasma progesterone concentrations (i.e. ≥ 3 ng/ml) on days 19-21, as well as the presence of a conceptus on ultrasonography (Prosound 2, Aloka Ltd., Japan) on days 30-45 followed by rectal palpation on day 60 post-artificial insemination (post-AI), were confirmed to be as pregnant. The cows which were not subjected to AI during their normal estrous cycle were considered as non-pregnant.

### 4.3. Feeding and management of experimental cows

All the experimental cows were fed and maintained as per the standard managemental practices followed at LRC, ICAR-NDRI, Karnal (Haryana) for KF cows. The concentrate mixture (CP=19.81% and TDN=70%) was composed of 33% maize, 21% groundnut cake (oiled), 12% mustard oil cake, 20% wheat bran, 11% deoiled rice bran, 2% mineral mixture and 1% common salt. Throughout the day, all of the animals had access to fresh tap water. The experimental animals were raised in a loose housing system with brick flooring and an asbestos roof and were free to move inside the controlled open area.

### 4.4. Blood Sampling

Blood samples (10 mL) from pregnant and non-pregnant cows were drawn aseptically from the jugular vein of each animal and placed in heparinized vacutainer tubes (Vacuette®-Na-heparin, Greiner Bio-One GmbH, Austria). The samples were taken on day 45 post-AI from the pregnant cows and day 0 (day of estrous on which animals were not subjected to AI) from the non-pregnant cows. Within 1 hour of collection, the samples were immediately transported to the laboratory under refrigerated conditions (icebox) and processed for automated blood cell count, plasma separation and isolation of neutrophil and PBMCs.

### 4.5. Determination of blood cell count using an automatic blood cell counter

100 μL of whole blood was taken in a 1.5 mL microcentrifuge tube (Tarsons) followed by automated analysis of blood count, as described in the manufacturer’s operational guidelines (MS4Se Automatic Blood Cell Counter, Melet Schloesing Laboratoires, Osny France). Blood cell count was performed to check the health status of the animals.

### 4. Isolation of plasma from whole blood

Freshly collected blood samples were centrifuged in 15 mL polypropylene falcon tubes (Tarsons) @ 1200 X g for 25 min to separate the plasma which was stored in storage vials at - 20 °C for the analysis of different pro-inflammatory and anti-inflammatory cytokines.

### 4.7. Quantification of pro-inflammatory and anti-inflammatory cytokines from the blood plasma

Plasma levels of anti-inflammatory (IL-4 and IL-10) and pro-inflammatory cytokines (IFNγ and TNFα) were estimated using bovine-specific ELISA Test Kits according to the manufacturer’s protocols. The optical density (OD) was measured by an ELISA reader (Multiskan Go, Thermo Scientific, Finland).

#### 4.7.1. Quantitative sandwich enzyme immunoassay for the determination of plasma interleukin-4 and interleukin-10 (anti-inflammatory cytokines)

Plasma IL-4 and IL-10 were estimated by “Bovine specific ELISA Kits” (Bioassay Technology Laboratory, Shanghai, China). The minimum detectable dose of bovine IL-4 (Cat. No E0036Bo) was 0.54 pg/ml, and the detection range of the assay was 1 pg/ml-380 pg/ml. The minimum detectable limit of bovine IL-10 (Cat.No. E0252Bo) was 2.52 pg/ml, and the detection range of the assay was 5-2000 pg/ml. The intra and inter-assay CV of both IL-4 and IL-10 were 8% and 10%, respectively.

#### 4.7.2. Quantitative sandwich enzyme immunoassay for the determination of plasma IFNγ and TNFα (pro-inflammatory cytokines)

Plasma IFNγ and TNFα were estimated by “Bovine specific ELISA Kits” (Bioassay Technology Laboratory, Shanghai, China). The minimum detectable dose of bovine IFNγ (Cat.No E0005Bo) was 2.35 pg/ml, and the detection range of the assay was 5 pg/ml-2000 pg/ml. The minimum detectable limit of bovine TNFα (Cat.No E0019Bo) was 5.56 pg/ml, and the detection range of the assay was 10-3000 pg/ml. The intra and inter-assay CV of both IFNγ and TNFα were 8% and 10%, respectively.

### 4.8. Isolation of neutrophils and PBMCs from whole blood

The isolation of neutrophils and PBMCs from whole blood was performed by the density gradient centrifugation method using Histopaque 1119 and 1077 (Sigma Aldrich, St. Louis, MO, USA) as described by Mohapatra et al., 2020. PBMCs were separated as a layer of the ring just above the Histopaque 1077 solution and neutrophils were separated at the interface of the Histopaque 1119 and Histopaque 1077 layers. The neutrophils and PBMCs were carefully separated to avoid mixing and washed with doubled volume of cold Dulbecco’s phosphate-buffered saline (DPBS, Himedia, India Pvt. Ltd) before being suspended in Dulbecco’s minimal essential medium (DMEM) (Sigma Aldrich, St. Louis, MO, USA) for further processing.

The number and viability of neutrophils and PBMCs were assessed by Haemocytometer (Reinfeld, Germany) using 0.4 % Trypan blue (Sigma, St. Louis, MO, USA) method as depicted by Alhussien and Dang, 2018. The viability of blood neutrophils and PBMCs was greater than 90% within the first few hours of the cell processing and declined gradually afterward. The purity of the neutrophils and PBMCs population was greater than 95% as evaluated by staining with pure May–Grunwald (HiMedia Laboratories, Pennsylvania, USA) for 2 min and Giemsa solution (HiMedia Laboratories, Pennsylvania, USA) for 20s and observed under oil immersion at 100 X (Olympus IX51 microscope).

### 4.9. Preparation of media for neutrophil culture

Chemically defined Dulbecco’s minimal essential medium (DMEM) was used to suspend isolated neutrophils. DMEM (Sigma, D2906-1L, without NaHCO_3 &_ Phenol red) was dissolved in sterile water (15.6g/l) and supplemented with sodium bicarbonate (1.2g/l). The pH of the resulting solution was adjusted to 7.3 using 1N NaOH and filtered through 0.22 μm Millex – GV filter unit (Millipore). The medium was aliquoted in sterilized 100 ml reagent bottles and stored at 4 ºC in the refrigerator.

### 4.10. Evaluation of key neutrophil functions

The functionality of neutrophils in the test animals was assessed by chemotaxis (velocity, distance, displacement and directionality), measuring activity of marker enzymes (myeloperoxidase and β-D glucuronidase) and nitric oxide production.

After isolation of neutrophils, one aliquot was used to determine chemotaxis and myeloperoxidase activity of neutrophil functionality and another aliquot was used for stimulation with or without LPS (1 μg/ml) for nitric oxide assay.

#### 4.10.1. Neutrophil Chemotaxis Assay

Chemotaxis assay was performed using the Dunn Chemotaxis Chamber (Hawksley DCC100) as per the manufacturer’s protocol. Neutrophils (suspended at 1 × 10^6^ cells/ml) were allowed to adhere to the coverslip coated with 7.5% culture-tested bovine serum albumin (Sigma-Aldrich, UK) for 20 minutes at room temperature. Initially, both annular wells were pre-filled with control medium (DMEM) and the coverslip seeded with cells was inverted onto the chamber in an offset position to leave a narrow filling slit at one edge for access to the outer well. The outer well was then drained and refilled with DMEM containing formyl methionyl-leucyl-phenylalanine (fMLP, 50μM) chemoattractant.

Cells were monitored under a microscope (Olympus, Tokyo, Japan) with a 40X phase contrast objective lens. The videotape recording was performed using a time-lapse option and individual frames were recorded at 60s intervals for 90 minutes using Q-CapturePro7 software. The videotape recording was converted to digital imaging for analysis by image J software (Java. Inc). Parameters like migration distance, speed, velocity & directionality were analyzed.

#### 4.10.2. Neutrophil Myeloperoxidase (MPO) assay

Myeloperoxidase (MPO) activity was measured, as reported by Bradley et al. (1982). Briefly, neutrophils (1 × 10^6^ cells/ml) suspended in DMEM was mixed with 9 volumes of 50 mM phosphate buffer (pH 6.0) containing 0.5% Cetyl trimethylammonium bromide (CTAB) and freezed at -20°C overnight. Samples were thawed the next day at 4°C followed by sonication in an ice bath. The samples were freeze-thawed three times followed by sonication for 10 s. The suspension was centrifuged (12000×g for 15 min) and the cell lysate was analyzed for MPO assay. All reagents and samples were brought to 25°C before performing the MPO assay.

The supernatant (0.5 ml) was mixed with 2.5 ml of the reaction mixture (50 mM potassium phosphate buffer, pH 6.0 containing 0.167 mg/ml of o-dianisidine dihydrochloride and 0.0005% H_2_O_2_). MPO degrades H_2_O_2_ to release oxygen free radicals which oxidize o-dianisidine dihydrochloride. The breakdown of H_2_O_2_ is directly proportional to oxidation of o-dianisidine dihydrochloride which is measured at 460 nm using a UVD-3500 double beam UV/visible spectrophotometer (Labomed, USA). The concentration of oxidized o-dianisidine dihydrochloride was calculated from its molar extinction coefficient (11.3 mM^-1^ cm^-1^). One unit of enzyme activity was defined as that oxidizing 1 μmole of o-dianisidine per min at 25°C and expressed as units/10^6^ neutrophils and calculated as follows:

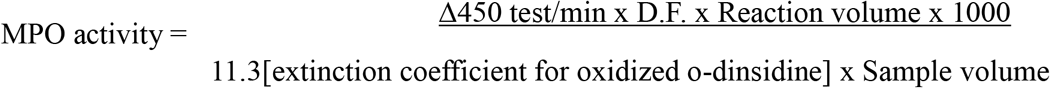

#### 4.10.3. Neutrophil β-glucuronidase assay

β-glucuronidase activity was determined according to the method of Stossel (1980). Briefly, 0.50 ml of neutrophil culture supernatant, 0.25 ml of PNPG and 0.25 ml of acetate buffer was added in clean and dried test tubes. The test tubes were then incubated for 5 h at 37°C. After incubation, the reaction was stopped by adding 1 ml of 0.1 M NaOH. The absorbance was determined at 410 nm using a spectrophotometer (LABINDIA® UV-VIS Spectrophotometer 3092) and the amount of PNP released was determined from the standard curve prepared with PNP. One unit of enzyme activity was defined as one millimolar of PNP liberated from the substrate per minute per 10^6^ cells.

#### 4.10.4. Neutrophil Nitric oxide production assay

Total nitric oxide production in neutrophil culture supernatant was determined by spectrophotometric measurement of its stable decomposition product i.e., nitrite (NO_2-_) using the protocol of Kasai et al., (1995). Briefly, the neutrophils suspended in DMEM was cultured in the presence or absence of LPS (1 μg/ml) for 3 hr in a humidified, 5% CO_2_ incubator. The supernatant of stimulated and unstimulated neutrophils was analyzed for nitric oxide according to the method of Kasai et al., (1995). Briefly, 50 μl of neutrophil culture supernatant and 50 μl of Griess reagent were added in a 96 well culture plate and incubated at 37°C for 45 min. The absorbance was measured at 550 nm and the concentration of nitrite (an indicator of nitric oxide) was calculated from the standard curve generated from serial dilutions of NaNO_2_, (0.125 μM-2.5 μM).

### 4.11. Evaluation of PBMCs functionality

The production of cytokines by the PBMCs is regarded among their key functional parameters which has been determined through relative mRNA expression of canonical pro-inflammatory and anti-inflammatory genes.

#### 4.11.1. Total RNA extraction from PBMCs

PBMCs isolated from each blood sample were adjusted to 1 × 10^6^ live cells/mL for carrying out expression studies. The RNA was extracted and purified by TRIzol reagent (Invitrogen, Carlsbad, CA) as prescribed by the manufacturer’s protocol. Briefly, cells were lysed by adding 1 mL of TRIzol reagent and the cell lysate was passed several times through a pipette followed by vortex for 5 min at room temperature (RT). The samples were incubated for 5 min at RT, followed by the addition of 0.2 mL of chloroform (HiMedia Laboratories, Pennsylvania, USA) per 1 mL of TRIzol and centrifuged at 12,000 × g for 15 min at 2−8 °C. The aqueous phase was transferred to a new tube and 0.6 mL of isopropyl alcohol (Sigma Aldrich, Darmstadt, Germany) was added per mL TRIzol, mixed and incubated for 10 min at RT. The samples were centrifuged at 12,000 × g for 10 min at 2−8 °C and the RNA pellets were washed with 1 mL of 75% ethanol, vortexed and centrifuged at 8500 × g for 5 min at 2−8 °C. The ethanol was removed and the pellets were left to air dry for 15 min. The dried RNA pellet was dissolved in 20 μl nuclease-free water (Thermo Scientific, USA) and its concentration was determined by measuring absorbance at 260 nm. The purity of RNA was evaluated by Biospec-nano Spectrophotometer (Shimadzu Corp., Japan) and judged by OD ratio at 260:280 nm. A ratio of (1.8−2.0) was considered for further processing. The integrity of each RNA sample was evaluated by agarose gel (2%) electrophoresis in Tris-EDTA buffer (0.002M EDTA) with (0.5 mg/mL) of ethidium bromide. Two intact bands of 28s and 18s without smearing indicated good quality and intactness of RNA. The RNA was then stored at −80 °C until further use.

#### 4.11.2. cDNA synthesis and real-time PCR (qPCR)

The extracted and purified RNA was subjected to DNase treatment by DNase1, RNase free (Thermo Scientific, USA) as prescribed by the manufacturer’s protocol. Total 500 ng of RNA from each sample was transcribed into complementary DNA (cDNA) by using Thermo Scientific Revert Aid First Strand cDNA synthesis kit (Thermo Scientific, USA). The RT reaction was carried out at 65 °C for 5 min, 42 °C for 50 min and 85 °C for 5 min in a thermal cycler (Bio-Rad, USA). Synthesized cDNA was kept at −80 °C till further use. Quantitative real-time PCR (qPCR) (Roche’s Lightcycler 480) was performed using Thermo Scientific Maxima SYBR Green qPCR Master Mix kit (Thermo Scientific, USA) according to the manufacturer’s instructions. Briefly, the reaction mix prepared was: 1 μl template; 5 μl (2×) SYBR green mixes, 0.5 μl each of reverse and forward primer, and 3 μl nuclease-free PCR grade water. Details of primers used for specific bovine IDO1, IL-10, IL-4, TGFβ1, IFNγ and TNFα genes have been shown in Table 1 (Sigma Chemicals Co., St. Louis, Missouri, USA). The following protocol and program were used: initial denaturation at 95 °C for 5 min, followed by 35 cycles at 95 °C for 30 s, 58 °C for 30 s, and 72 °C for 30 s and then 72 °C for 5 min and final holding temperature was 4 °C. The expression of each gene was analyzed in triplicate and normalized with β-actin and GAPDH which were used as the housekeeping gene. The relative gene expression level was evaluated by the 2−ΔΔCt method.

**Table 1.**
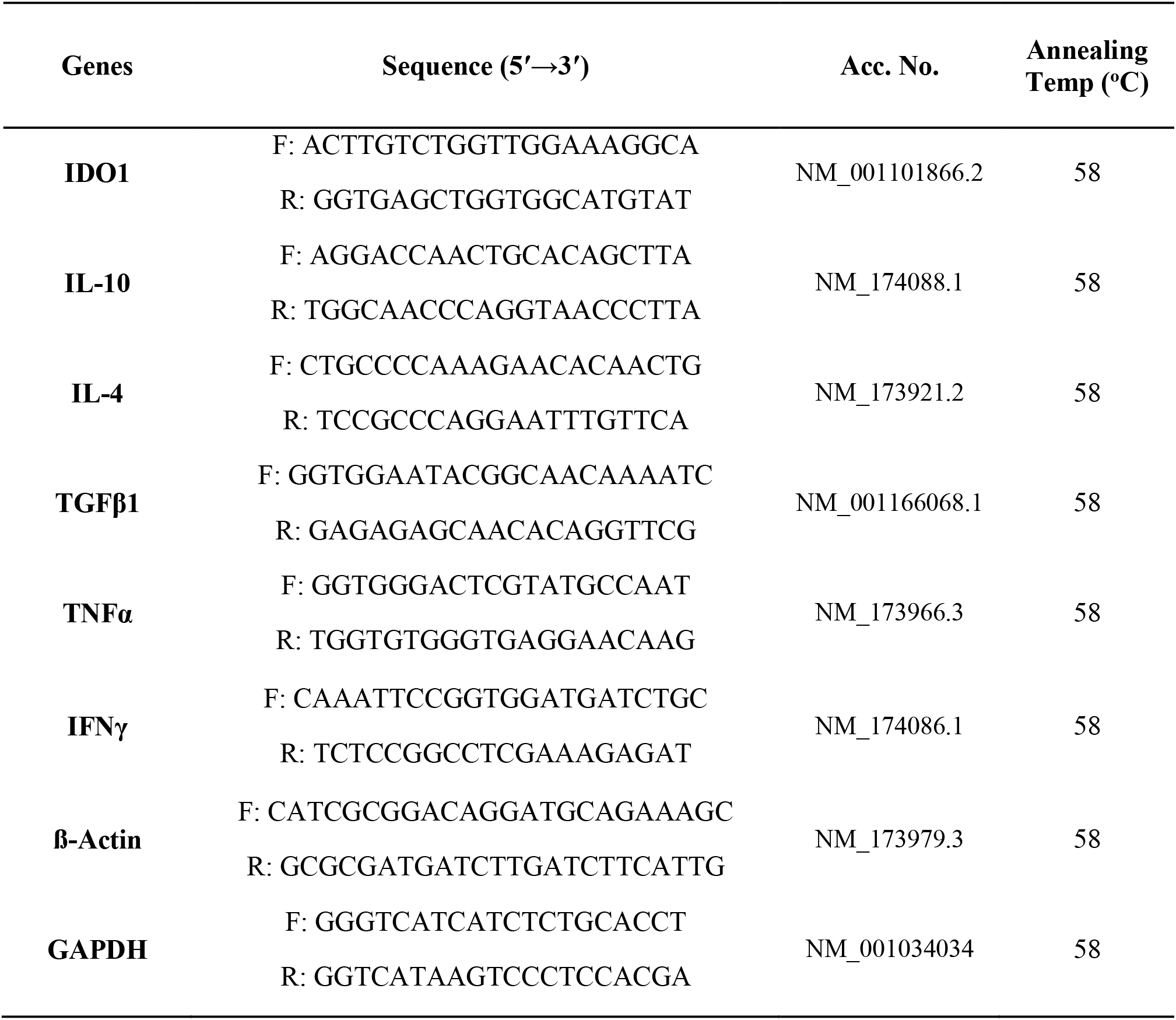
Details of primers used in the study.

### 4.12. Statistical analysis

Statistical analysis was performed using the GraphPad Prism software (version 8.0.2, Graphpad Software, San Diego, CA, USA). All the data obtained from the study were expressed as mean ± standard error of the mean (S.E.M.) and were analyzed by two-way analysis of variance (ANOVA) followed by Tukey’s multiple comparison tests. A minimum of three independent experiments was performed for each experimental condition tested. The value of P<0.05 was considered to be statistically significant.

## 5. Conclusions

The present study highlights for the first time an insight into the possible role of IDO1 in modulating the functionality of neutrophils and PBMCs as well as regulating the maternal systemic cytokine balance/shift during early pregnancy establishment in bovines. Upregulation in IDO1 expression during pregnancy might be crucial to modulate or rather suppress the functionality of neutrophils in terms of chemotaxis, enzymes’ activity (myeloperoxidase and β-D glucuronidase) and nitric oxide production which are considered as necessary to eliminate the foreign antigen. Also, increased IDO1 expression might be necessary to modulate the lymphocyte functionality so as to produce and maintain a favorable cytokine milieu in order to prevent rejection of the semi allogeneic fetus. We believe that extensive research on the development of IDO1 as biomarker may be carried out in future to study its possible role in improving pregnancy outcome in the bovines.

## Acknowledgements

The authors would like to thank the Director of the Institute for providing all the facilities to carry out this research work.

## Author contributions

Conceptualization: S.K.M.; Methodology: S.K.M., D.C..; Validation: S.K.M., D.C.; Formal analysis: S.K.M.; Investigation: S.K.M., R.K., A.K.D; Resources: A.K.D.; Writing -original draft: S.K.M., B.S.K.P.; Writing - review & editing: R.K., A.K.D.; Visualization: S.K.M.; Supervision: A.K.D.; Project administration: A.K.D.; Funding acquisition: A.K.D.

---

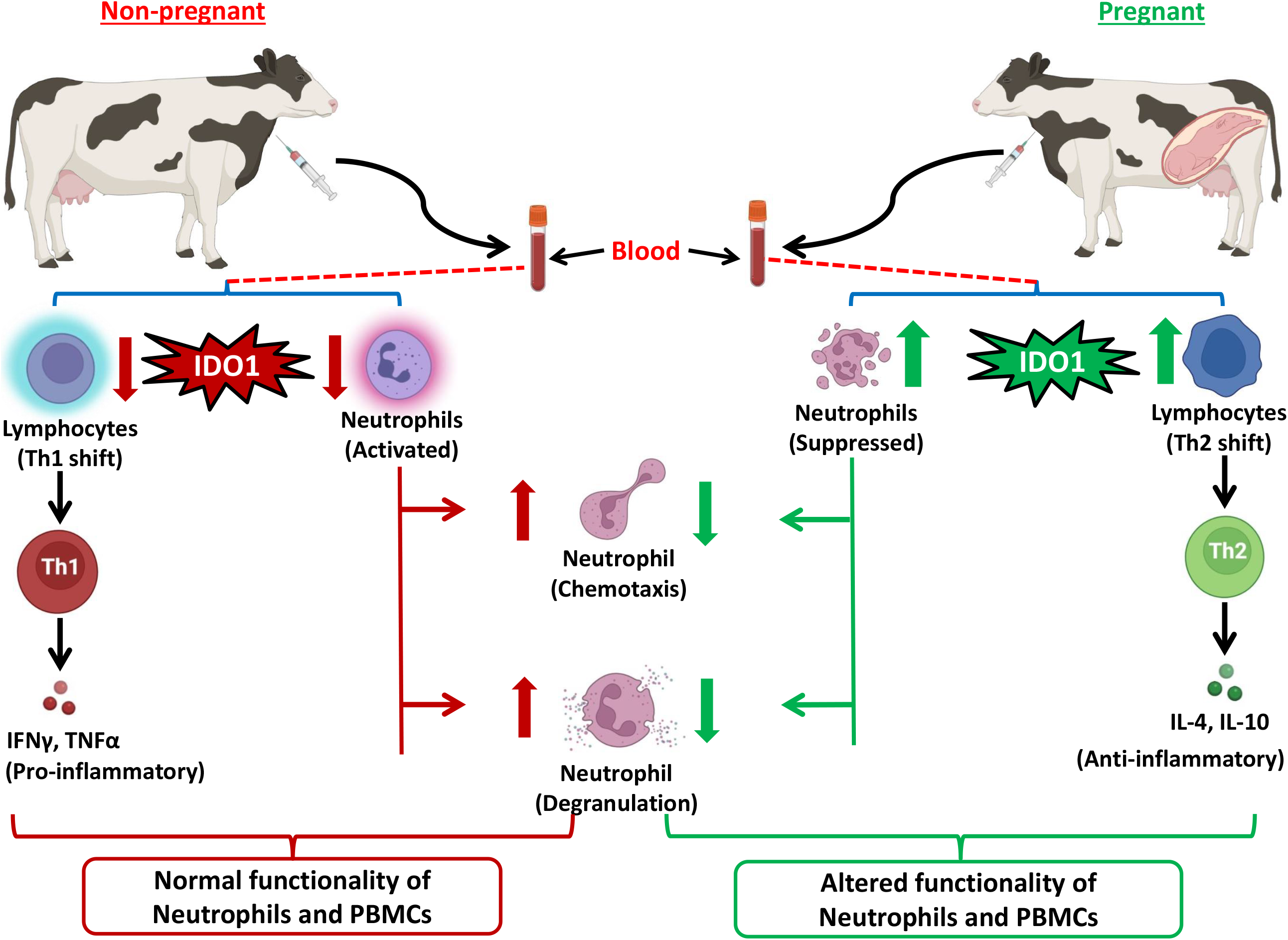

